# A novel two-indenter based micro-pump for lab-on-a-chip application: modeling and characterizing flows for a non-Newtonian fluid

**DOI:** 10.1101/2020.02.15.951111

**Authors:** Ravi Kant Avvari

## Abstract

Inspired by the feeding mechanisms of a nematode, a novel two-indenter (2I) micro-pump is analyzed theoretically for transport and mixing of a non-Newtonian fluid for the purpose of lab-on-a-chip applications. Considering that the viscous forces dominate the flows in microscopic regime, the concept lubrication theory was adopted to device the two-dimensional flow model of the problem. By approximating the movements of the indenter as a sinusoidal function, the details of the flow were investigated for variations in – frequency of contraction of the first value keeping the second valve at higher occlusion, and occlusion. The study indicates that occlusive nature of the second valve leads to the large pressure barrier which prevents the fluid to enter into the neighboring compartment. Transport occurs as the lumen opens to develop a suction pressure. Pressure barrier is found to be highest for dilatants followed by Newtonian and pseudo-plastics. Shear stress dependency on frequency the contraction of the first value is highest for lower values of flow behavior index. In conclusion, the study provides details connecting the flows resulting from the indentation of the front-end indenter to the frequency of indentation, geometry and rheology of the fluid, thus facilitating optimal design of the micro-pumps.

## 1. Introduction

A micro-pump is a miniaturized device that allows for pumping fluid samples such as drugs (drug delivery) and other compounds/ analytes to the target region with precision. They form an integral part of the Micro Drug Delivery System (MDDS), a miniaturized device that uses a micro-needle, micro-diaphragm, micro-valve and micro-pump actuated by various means such as – pneumatic, piezo, electrostatic, electroosmotic and electrochemical [1–3]. They also serve to perform various microfluidic related operations such as the sampling, filtration, dilution, reaction, mixing, and biosensing [4, 5]. Because of such diverse applications, they are readily used as an integral component of the Micro Total Analysis Systems (μTAS) [6], Lab-on-a-Chip and Point of Care (POC) testing systems [7] that caters to various applications such as drug delivery and pharmaceutical applications [8]. Measurement of flow and pressure is central to the design of micro pumps for such applications. In analytical systems, the micro-pumps transfer a fixed amount of reagent to another chamber to participate in the reaction process which processes such as transport, mixing, separation and sensing for analyte concentration. Systems such as the μTAS require controlling and transferring of tiny volume of fluids to perform functions of reactions and analysis of the solutions [9, 10]. Recent studies involving a more sophisticated design have developed lab-on-chip models of the gut to study the physiology of human intestinal cells and immune cells in relation to their interaction with the flow [11, 12].

Since the length scale of channel is of the order of micrometer, the transport of the fluids is dominated by the viscous forces and the inertia has negligible role in transport. Considering that the flow in the micro pump is driven by wall momentum; where, as the wall undergoes radial movement the fluid particles beneath the wall also move in accordance to obey the law of momentum conservation. The velocity of fluid particles at the wall equals the wall velocity due to the no-slip boundary condition; however, as we move towards the centre of the lumen, the velocity reduces. The mechanical actuation of the wall have been demonstrated in numerous systems that are driven by means of pneumatic [13–15], electrostatic [16, 17], electromagnetic [18, 19], or piezoelectric [20, 21] actuators. Diaphragms that can be actuated electrically, using two control valves have been used to regulate the flow of fluids in the channels. Micro-pumps using three diaphragms are used in series to perform the complex task of peristaltic-like transport in the channels. As the diaphragm is moved (up/downward either periodically or non-periodically), the flows are developed in accordance to the wall motion. At microscopic scale, where the inertial forces are low in comparison to viscous forces (assuming low flow rates resulting in Re≪), such pumps transfer their mechanical energy to help push the fluid along the direction of motion to cause the flow [2, 22]. In other words, diaphragm acts as a source of mechanical forces to induce peristalsis in the channel through generation of pressure forces. Peristaltic transport in biological systems is generated by a progressive wave of contraction/expansion of the wall musculature which helps in mixing and transferring the contents to the lower parts of the tube [23–25]. Due to challenges in replicating such systems in laboratory, micro fluidic systems comprising of a number of pumping stages have been employed for developing such peristalsis like motion in engineering applications. Authors have developed systems containing three chambers arranged in a series [26, 27] that are actuated by the piezoelectric disk to cause the opening or closing of the chamber. Such systems have been applied for drug delivery and various biological applications such as to perform the PCR (polymerase chain reaction) [28, 29]. A two-stage discrete peristaltic pump by Berg et al. [30] showed that peristalsis-like motion can also be demonstrated by using two chambers only to develop static head pressure and flow rates that are comparable to the three-stage pumps. Suggesting, that the dynamics involved in the spatio-temporal changes of the fluid velocity and pressure is of importance in the design of micro-pump for various applications.

In this paper, we have analyzed a novel pump based on the feeding behavior of a nematode and performed a semi-analytical study of the two-indenter pump by approximating Navier–Stokes equations using lubrication theory. We employ the analytical results of the peristaltic transport provided by Shapiro et al. [31] and Misra et al. [32] for formulating the approximate solutions for two-indenter driven peristaltic pump.

## 2. Mathematical modelling

### 2.1 Design of a novel 2I micro-pump

The design of the micro-pump is inspired by the feeding behavior of the *Caenorhabditis elegans*, a free-living nematode that is 1mm in length. The feeding behavior is driven by the pharyngeal motion of the alimentary system. It is made up of 20 muscle cells that help in initiating the mechanical movements by undergoing contraction. These muscle cells forms various segments of the alimentary canal such as the corpus, isthmus, and the terminal bulb. The mechanical activity of these muscles or the pharyngeal behavior involves pharyngeal pumping and isthmus peristalsis [33]. The muscles contract in a coordination manner to facilitate the food entry in the following process. The process is initiated by the contraction of corpus together with the anterior isthmus which helps in creating a vacuum to cause inflow of the external fluid into the alimentary canal. The nematode uses special tricks in sorting the food (that is, bacteria) by allowing only the fluid to exit the channel; luminal closure triggered by the relaxation of corpus allows for compression of the chamber to drive the outflow of the fluid [34, 35].

The feeding behavior of the nematode is approximated using a two indenter pump, where we consider the first valve to drive inward and outward flow of the fluid by regulating the valve position. From mechanics point of view, we speculate that the closure of the valve pressurizes the chamber to cause outflow of the fluid. Since the second indenter is held stationary and at higher occlusion, the flow of the contents through the constriction is prevented as it does not allow for free flow. We also speculate that such a system would capture the dynamics of flow that occur in the nematode during the ingestion.

### 2.2 Mathematical model for the micro-pump

A schematic of the pump is shown in **Fig. 1** having two indenters, first and the second indenter, for developing the pressure forces to drive the flow. We consider transport in a micropump that is driven by the channel walls; especially by first indenter keeping the second indenter static. The geometric model of the peristalsis pump has a channel length of 10 units (dimensionless form), having the indenters placed at 4.5 and 8.5 units respectively. Since the nematode is characterized by having a long channel of higher aspects ratio, we assumed a radial dimension of 1 unit and aspects ratio of 1:10 (radial to length dimension). Since, for small length scales, where the flows are dominated by the axial dimension in comparison to the radial ones, we have adopted the lubrication theory [31] to develop the mathematical model. At such aspects ratio, an order of magnitude difference, the variation in the pressure across the radial direction is negligible to affect the flow, and the inertia of fluid has negligible contribution to the fluid momenta.

**FIG. 1A.**
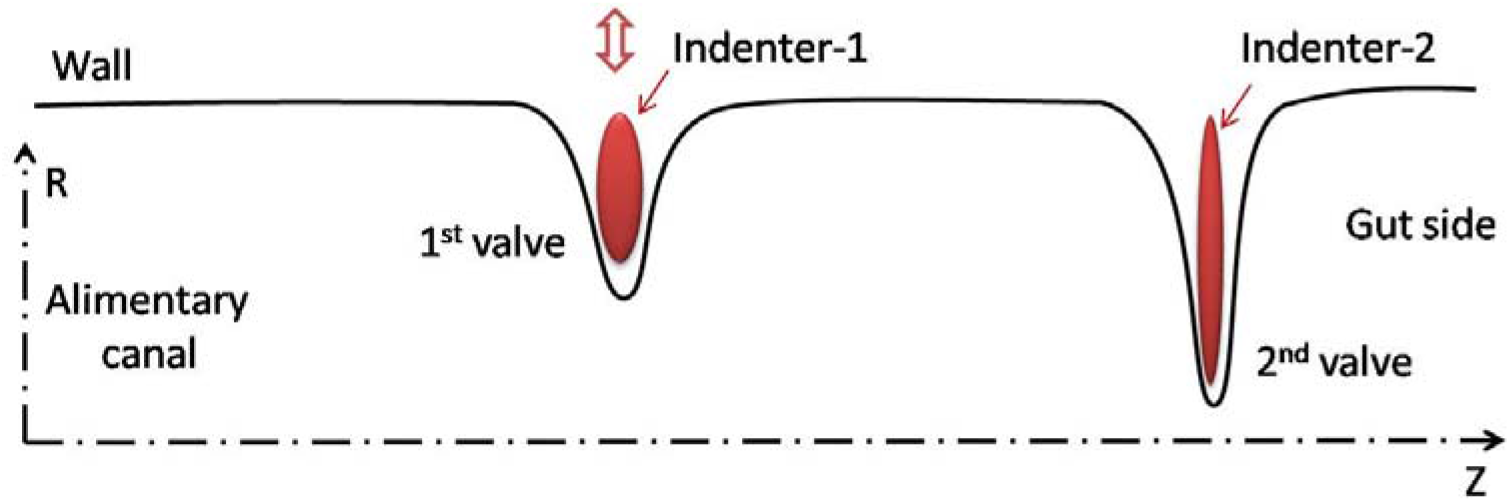
A two-indenter model of the micro-pump having one of its ends highly occluded by the indentor-2 to prevent flow of contents and indentor-1 as a source of stationary contraction to induce suction in the channel during opening and discharge during closure.

**FIG. 1B.**
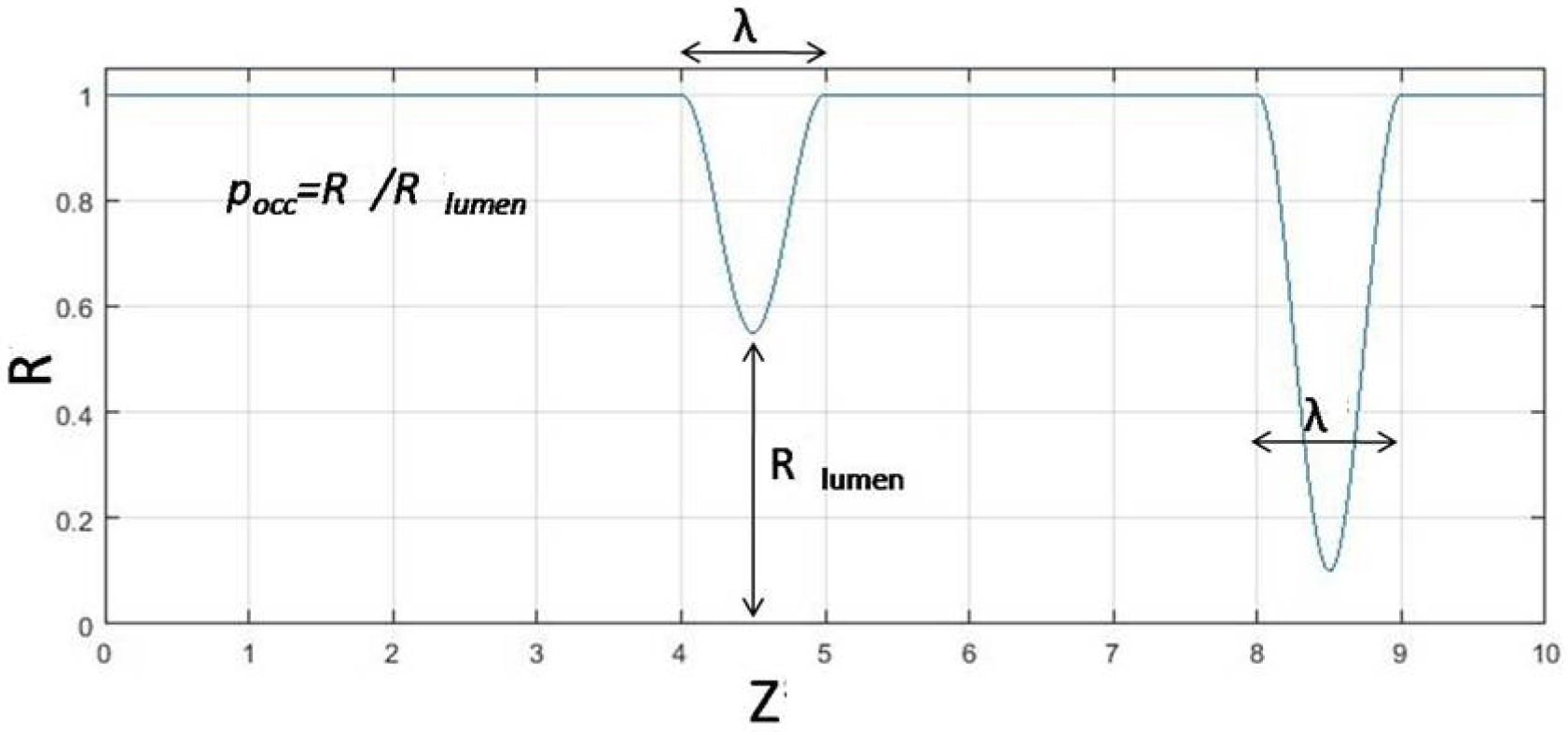
The mathematical model of the two-indenter micropump indicating the stationary wave due to the 1^st^ indenter (front-end) and an occluded tube due to the 2^nd^ indenter (rear-end) of certain wavelength.

In the present study, we have assumed the radial deformations of the wall (assumed to be non-flexible) as a stationary wave deformation modeled using a sinusoidal function of axial dimension ‘z’, time ‘t’, and having an angular frequency, ω=2πf, and an amplitude function R(z, t) as follows,

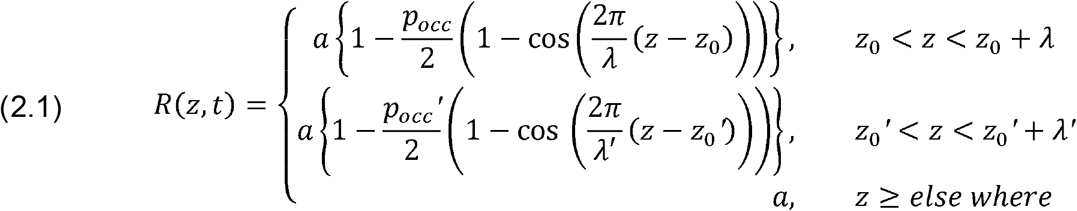

Where,

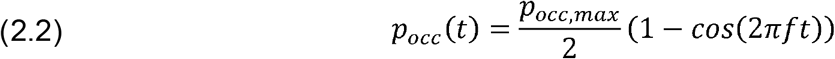

The equations governing the flows due to the stationary contraction and the constriction at the other end of the tube is solved semi-analytically using lubrication theory as derived by Meijing et al, adopted for non-Newtonian fluid [36, 37] and modified to approximate the flow due to the desired boundary conditions [24].

The equations governing the flows of an incompressible fluid of constant dynamic viscosity in cylindrical coordinate system (*r, θ, z*) are given by,

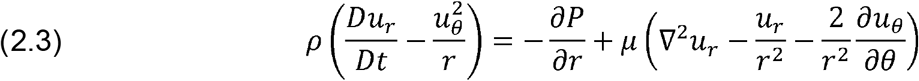

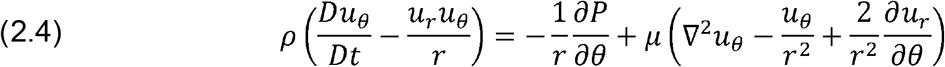

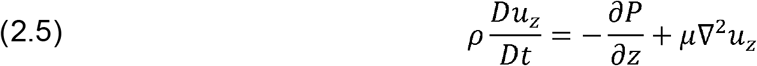

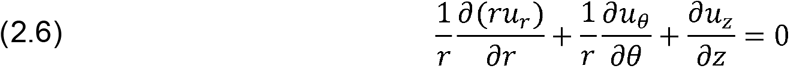

Where,

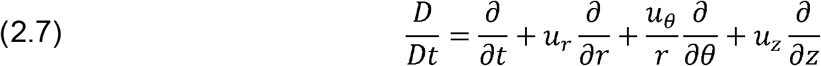

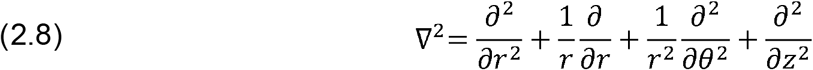

The fluid is modeled as power law,

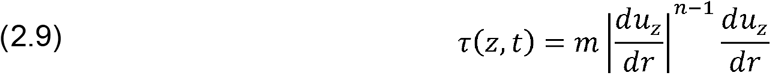

Where *m* is known as the flow consistency index and *n* is the flow behavior index. For values of *n*<1, the fluid responds as a pseudo plastic (e.g. milk), *n*=1 the fluid behaves like a Newtonian (e.g. water), and *n* > 1 the fluid acts like a dilatants (e.g. sugar solution, suspension of rice starch).

We define the following non-dimensional variables; where, *u_r_, u_z_* are the velocity components, *P* is the luminal pressures, *μ* is viscosity, and a is radius of the channel.

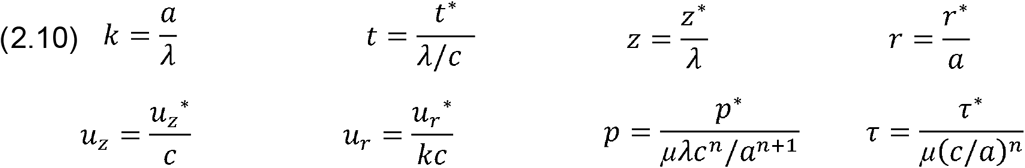

The wavelength (*λ*) of contraction due to the first indenter and channel length (a) were considered as the length scales and *λ/c* as the time scale for non-dimensionalization.

Navier Stokes equations in cylindrical coordinate system after approximation to lubrication limit (long wavelength approximation - *λ* ≫ *a*, resulting in *Re* ~ 0 or negligible inertial effect and dominated by viscous forces) is given by,

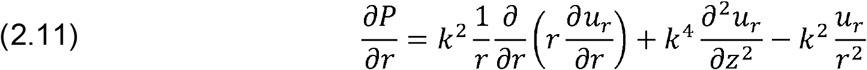

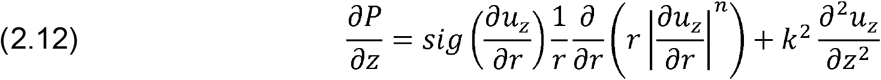

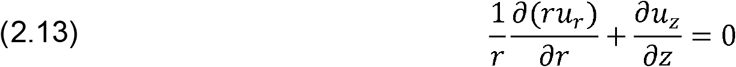

We apply boundary conditions (BC) to obtain desired flow of no-slip condition at the wall and subject to given pressure and velocity BC’s we obtain,

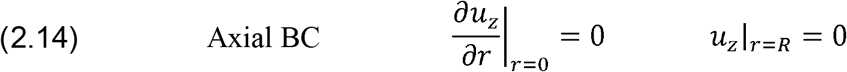

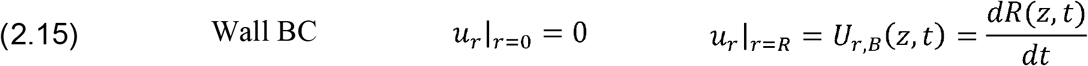

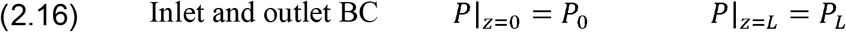

Using the approximate model of the flow, we derive the axial and radial velocities of the fluid is as,

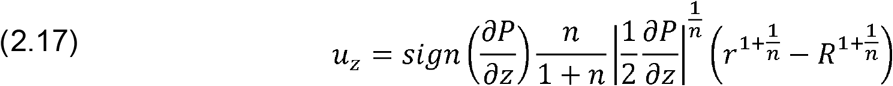

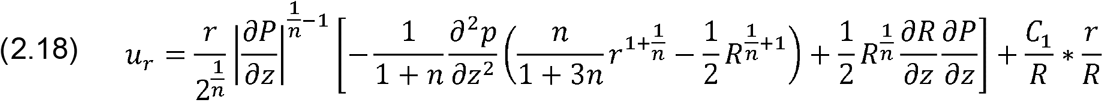

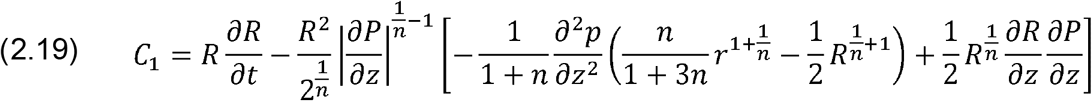

Integrating **Eq. (11, 12)**, the pressure gradient in the channel and the pressure field is obtained,

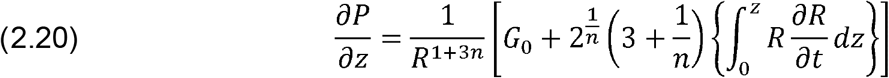

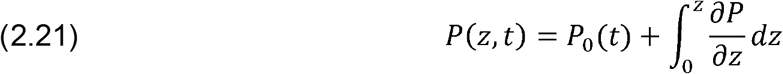

Using the definition of shear stress, the shearing effect at the wall is found as follows,

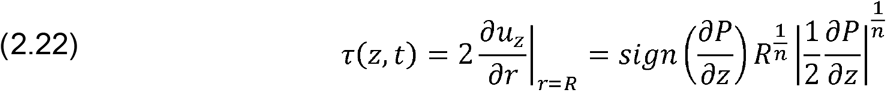

We also determine the instantaneous flow rate at a given location in the channel and the averaged flow rate by,

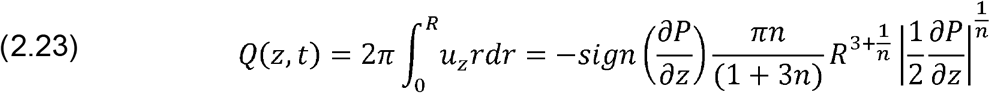

## 3. Experimental study

The mathematical model was solved on a MATLAB platform, by converting the governing equations into an algebraic form. The geometry was resolved using grid of size 60 (radial direction) and 600 (axial direction). The integral terms were discretized using Simpson’s rule and the differential terms using Runge-Kutta method (4^th^ order). Parameters of interest are frequency of oscillation of the front-end indenter (f), wavelength (λ), occlusion (*p__occ_*) and viscosity (*μ*). Simulations were performed by considering variation of two parameters (a 2 parametric study; where one parameter is flow behavior index) at a time to capture the peak pressure, flow rate and shear stress. Since the rate of flow occurs maximally at 3/4^th^ of the time period of oscillation of the front-end indenter, where the rate of closure of the wall is maximum, simulations were performed during this time point (*t*=3/4*T*).

## 4. Results and discussions

We have modeled the micro-pump by considering the indenter in such a way that it assumes a sinusoid impression on the wall. The indenter can be actuated by a motorized system which allows for the adjustment of the occlusion at the valves. By fixing the second value at a higher occlusion and allowing for stationary contraction at the first valve, we have simulated the dynamics of the non-Newtonian fluid transport both for closing (**Fig. 2**) and opening contraction (**Fig. 3**).

**FIG.2.**
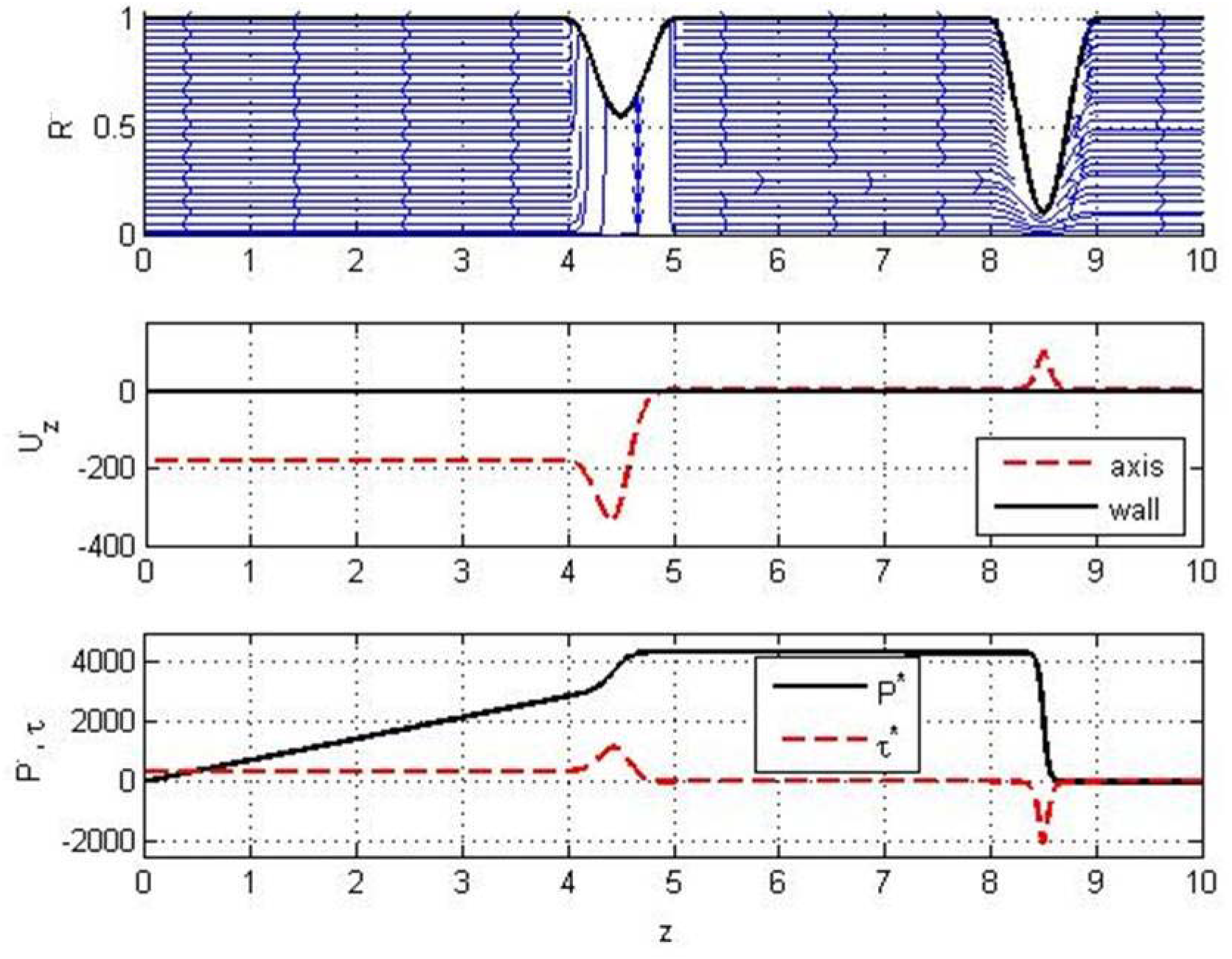
Flow due to stationary contraction (closing type) showing streamline patterns (top panel) in a channel for a wave of width = 0.10. Axial velocity is indicated in the mid panel, and pressure and shear stress in the bottom panel. Plots correspond to base parameters – n=0.5, R*=1mm, L*=20mm, μ=0.01Pa.s, c=1mm/sec, p_occ_max_ = 90%, λ*=2mm. Note: * indicates the dimensional form.

**FIG. 3.**
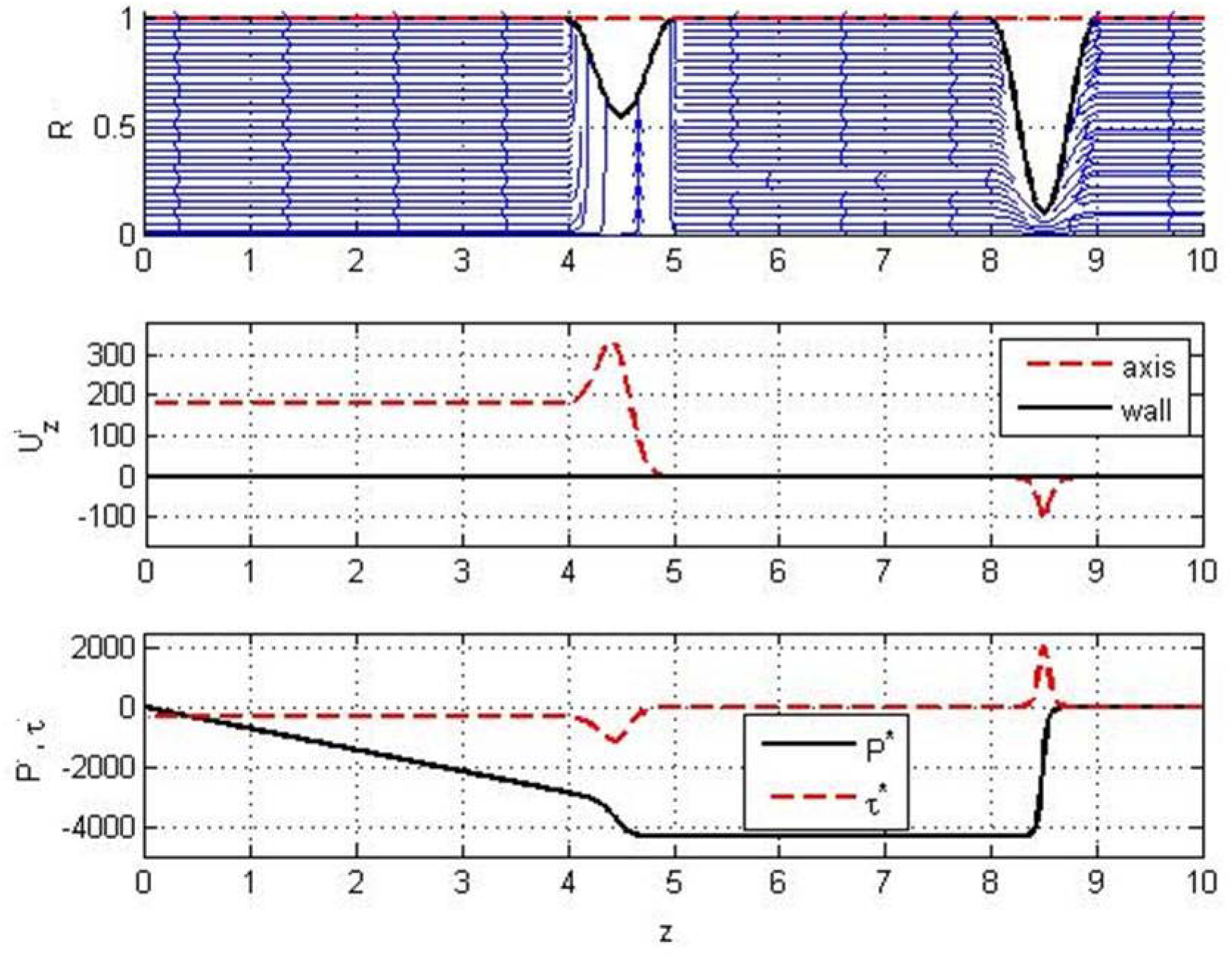
Flow due to stationary contraction (opening type) showing streamline patterns (top panel) in a channel for a wave of width = 0.10. Axial velocity is indicated in the mid panel, and pressure and shear stress in the bottom panel. Plots correspond to base parameters – n=0.5, R*=1mm, L*=20mm, f=6Hz, p_occ,max_ = 90%, λ* = 2mm, μ=0.01Pa.s.

The contraction at the second valve has been frozen to a fixed occlusion of 90% to prevent passive flow of contents between the compartments, which, as a consequence leads to generation of a high pressure barrier. By such simplification in the simulation study we have focused to capture the events of pressure and velocity changes for stationary contractions of various parameters such as frequency of contraction (rate of opening and closing), occlusion and wavelength. Closing of the first valve pushes the fluid in the lumen in the downward direction causing increase in the pressure at the region. However, due to the obstruction at the second valve the fluid in between the valves have to push strongly to escape through the narrow space at the second valve. The chamber formed is already at higher pressure due to compression which is indicated by a straight line in **Fig. 2**. The pressure drops drastically at the second valve and moderately at the first valve in a nonlinear manner. Since the first valve partly allows for fluid transport due to moderate occlusion, by virtue of higher pressure at the first valve, the fluid is pushed outward proportional to the pressure gradient (linear decrease in pressure). Due to higher pressure gradients the flow rates appear to be higher at both the valves with generation of high fluid stress. Direction of flow at the inlet is outward due to generation of negative pressure gradient.

Flow due to stationary contraction of opening type performs opposite to the closing type (**Fig. 3**). With the underlying principles remaining the same, the opening of the lumen creates suction at the first valve leading to inflow of the contents from outside (marked by a negative pressure gradient). Inflow of contents from the outlet is negligible due to the presence of highly occlusive valve.

The characteristic of fluid transport is apparently decided by the rate of closure of the first valve and depending on the nature of fluid, the flow details vary. As shown in **Fig. 4**, the nature of fluid can directly affect the fluid transport in the micro-pump. Variation of pressure over its length for various values of flow behavior index suggests that the magnitude of pressure generated is highest for the dilatants followed by Newtonian and pseudo-plastic fluids. Presuming that the flexibility of the wall has a negligible effect on the pumping of the fluid at a given rate, we identify that the forces required for pushing the dilatants are much higher than Newtonian fluids; thus lending it more difficult to pump. Due to higher pressure gradients at the second valve a relatively higher fluid stresses are developed in dilatants. Changes in the flowrates are found to be negligible for different fluids.

**FIG. 4.**
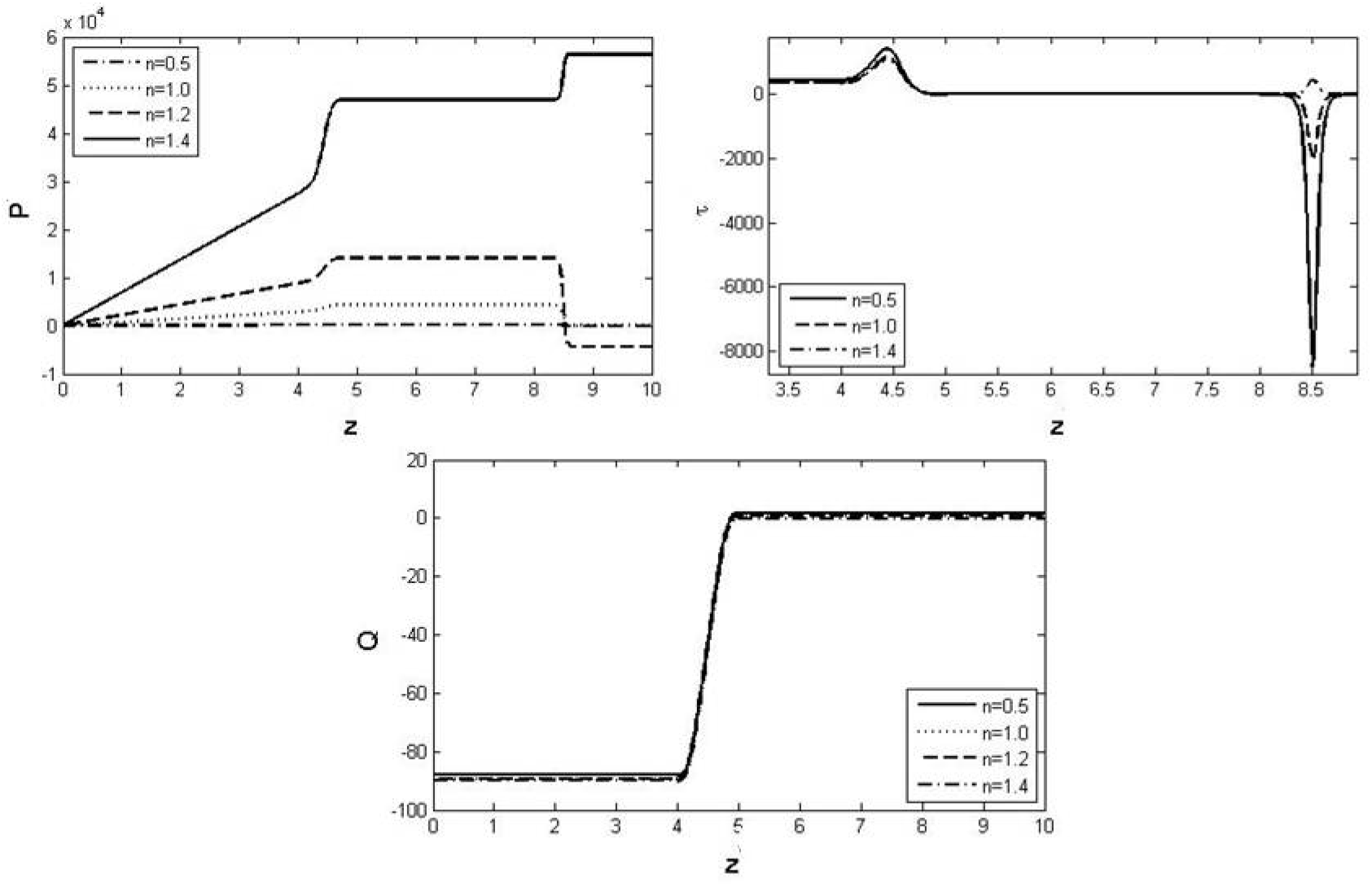
Effect of flow behavior index on the luminal flow at t=0.75Tsec (closing type).

Effect of maximal occlusion on the closing of the lumen is shown in **Fig. 5**. Spatial variation of the pressure is invariable and depends on the degree of occlusion. At higher occlusion the chamber formed in between the two valves are pressurized heavily compared to those of lower occlusion. As a result of this, higher flow rates are developed which has an effect of generating higher fluid stress.

**FIG. 5.**
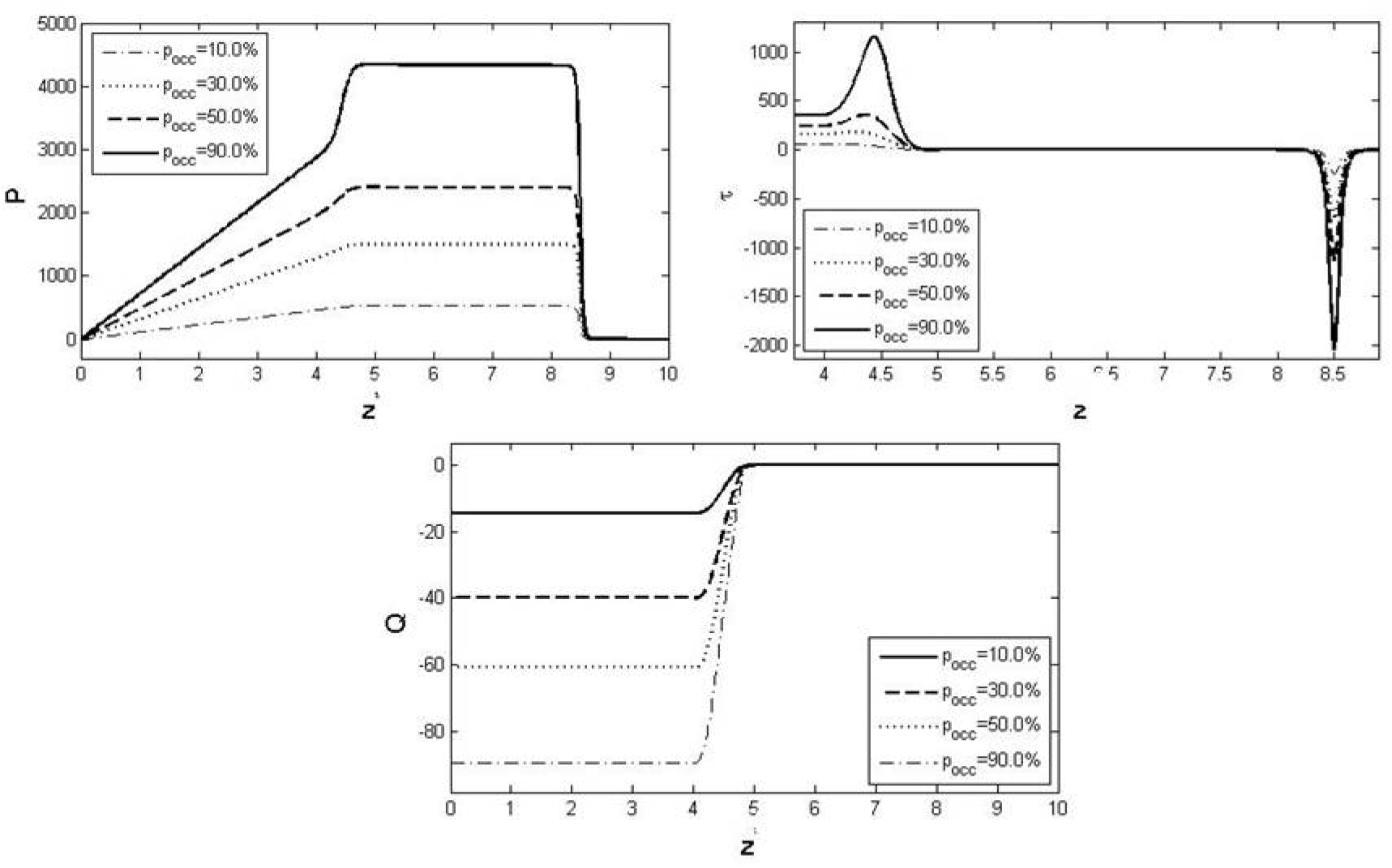
Effect of maximal occlusion on the luminal flow at t=0.75Tsec (closing type). The maximal occlusion is the peak occlusion which the contraction can process in a sinusoidal manner and approaches peak wall momentum at 1/4^th^ of the time period of the rate of occlusion.

Considering a frequency of 1-12Hz occurs in various biological systems (such as the small intestine) we have simulated flow at 1, 4, 9 and 12Hz (**Fig. 6**). Increase in the frequency of contraction has an effect of increasing the pressurization of the chamber. The relative increase in the pressure rise appears to be proportional. This increase in pressure proportionately affects the shear stress and the flow rates.

Effect of increasing the span of contraction (wavelength) from a 50% reduction in the baseline wavelength to a 200% increase, leads to generation of a relatively higher pressures in the lumen and an increase in the flowrate (**Fig. 7**). Increasing the width of contraction increases the span of pressure variation and the fluid stress; allowing more distributing stress over longer dimension of the pump.

**FIG. 6.**
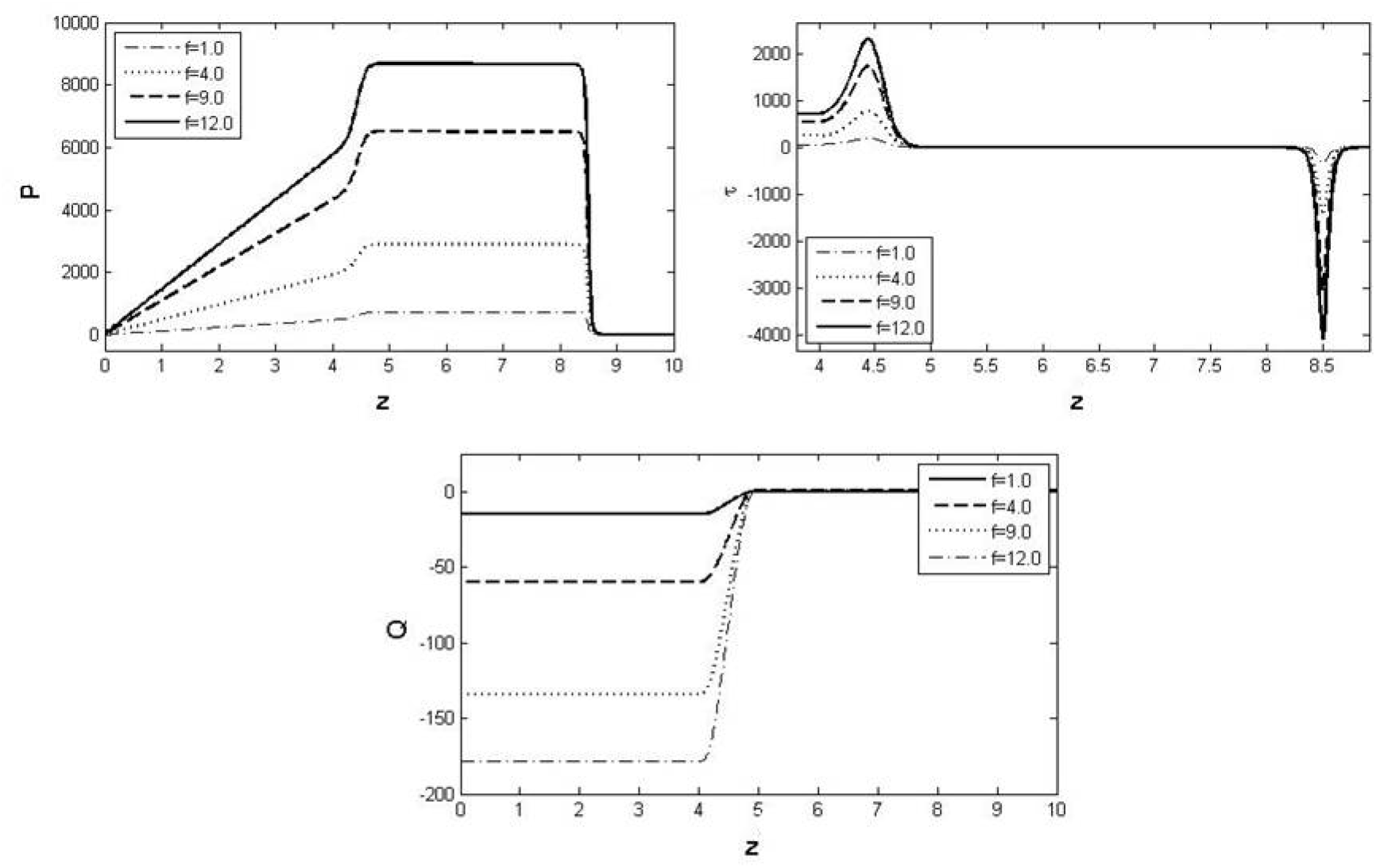
Effect of frequency on the luminal flow at t=0.75Tsec (closing type).

**FIG. 7.**
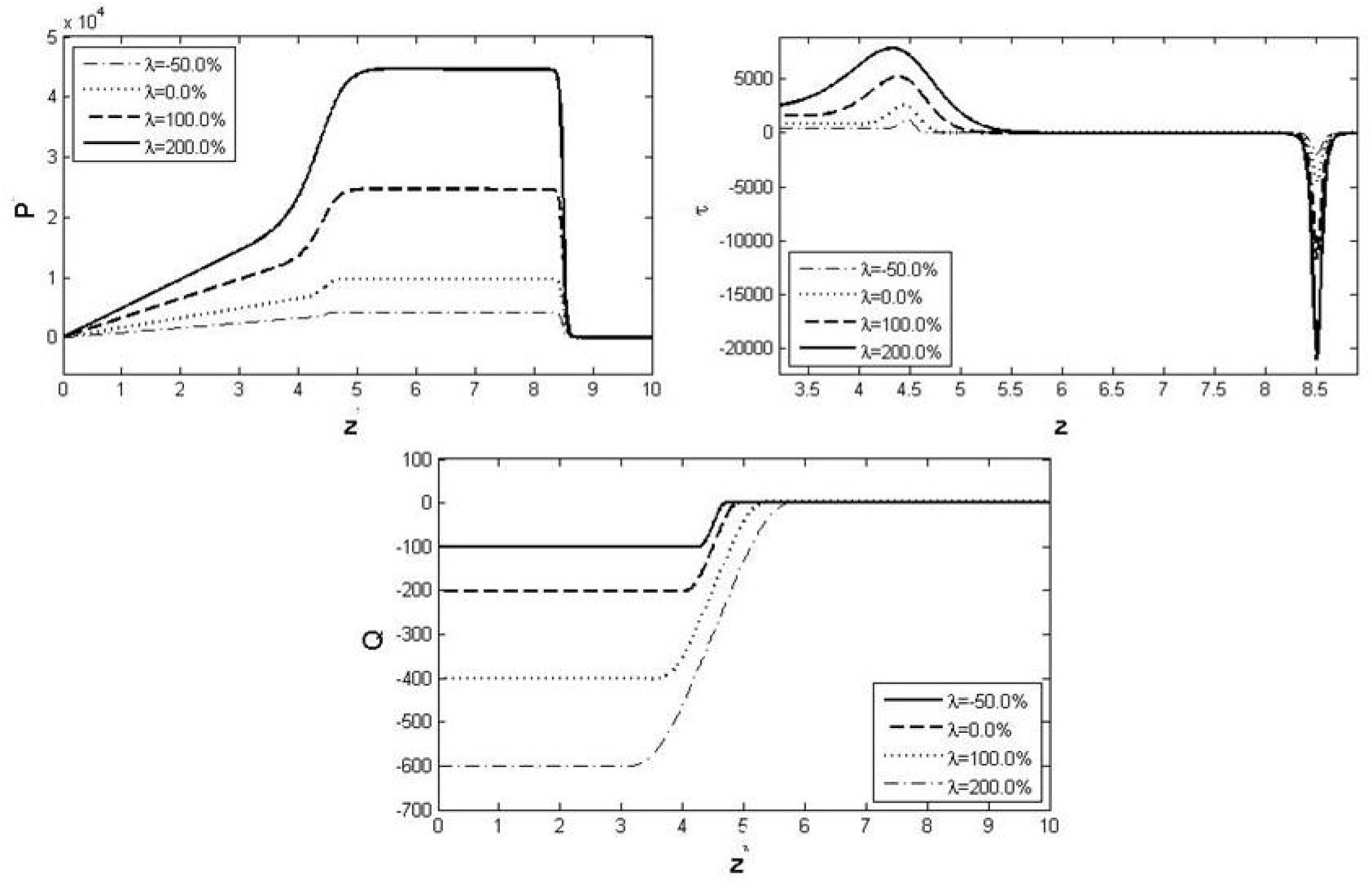
Effect of wavelength on the luminal flow at t=0.75Tsec (closing type).

To assess the performance of pumping across various parameters of the micro-pump (geometry and fluid), a parametric study was performed. Effects of occlusion on the flow are shown in **Fig. 8**. Pressure values at close to the inlet (L/6 units of distance from inlet) of the pump was considered for comparison, rather than considering the peak pressure generated since the pressure variation is linear and related to global flow rate. With increase in the occlusion of the 1^st^ valve, we observe a linear increase in the pressure at lower occlusion and reduction in the slope for higher occlusion; suggesting that the extent of pressure rise is not same at higher occlusion. Pressure values are highest for dilatants and lowest for the pseudo-plastic fluids. The pressure rise due increase in occlusion is highest for dilatants and decreases significantly with decrease in flow behavior index. Pseudo-plastic may appear to show negligible variation when compared to dilatants.

**FIG. 8.**
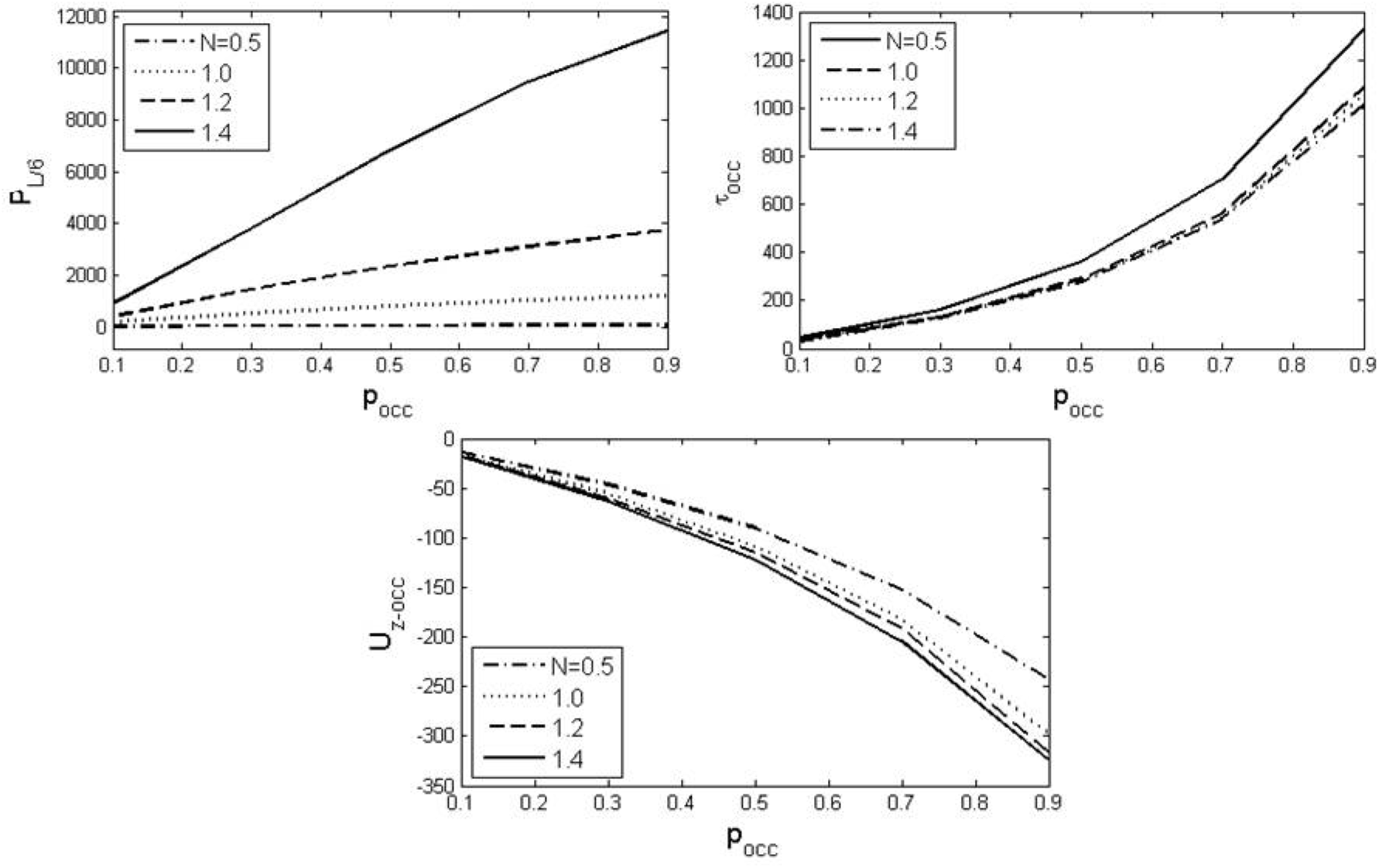
Effect of occlusion on the flow for fluid having flow behavior index = 0.5 (pseudo-plastic), 1.0 (Newtonian), 1.2 (dilatants), and 1.5 (dilatants).

Peak wall shear stress is found to be increasing nonlinearly with increase in occlusion; with negligible change for higher values of ‘n’ (**Fig. 8**). The peak axial velocity increases with increase in the occlusion of the lumen; with highest for dilatants and lowest for pseudo-plastic. Increasing frequency of contraction favors development of higher pressure for dilatants and to a least extent for pseudo-plastic fluid (**Fig. 9**). Shear stress dependency of frequency is highest for lower values of ‘n’ with higher slopes. Axial velocity of fluid is found to be linear related to frequency rise with highest slope for dilatants. Flow dependency on viscosity is the same, however, shifts with type of fluid considered (**Fig. 10**). Dilatants develop higher pressure forces and lower shear. Increase in the wavelength of the wave brings about a near linear change in pressure, shear and velocity (**Fig. 11**). For dilatants, the change is highest for the luminal pressure and axial velocity and lowest for shear stress developed at the wall.

**FIG. 9.**
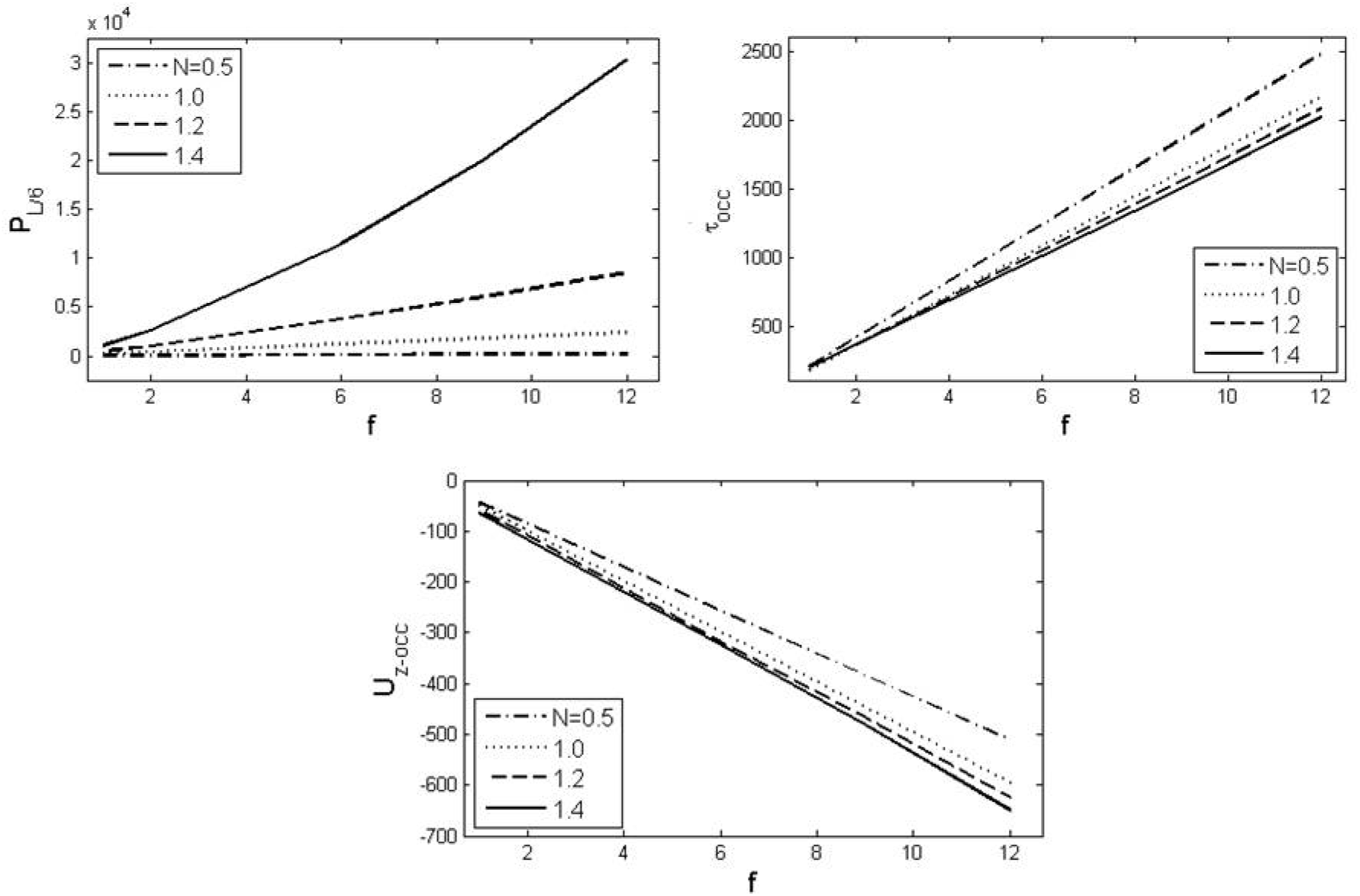
Effect of frequency on the flow for fluid having flow behavior index = 0.5 (pseudo-plastic), 1.0 (Newtonian), 1.2 (dilatants), and 1.5 (dilatants).

**FIG. 10.**
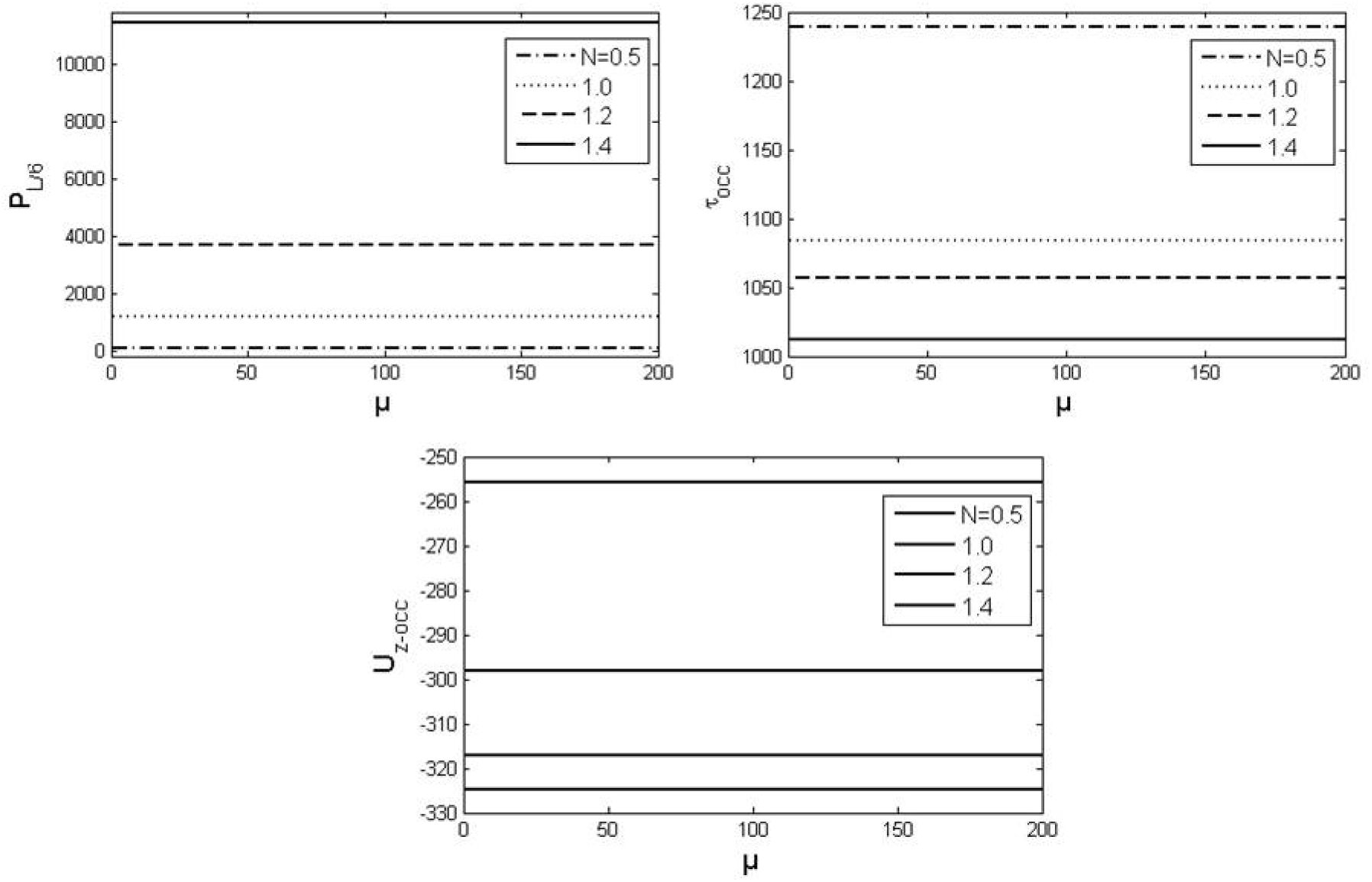
Effect of viscosity on the flow for fluid having flow behavior index = 0.5 (pseudo-plastic), 1.0 (Newtonian), 1.2 (dilatants), and 1.5 (dilatants).

**FIG. 11.**
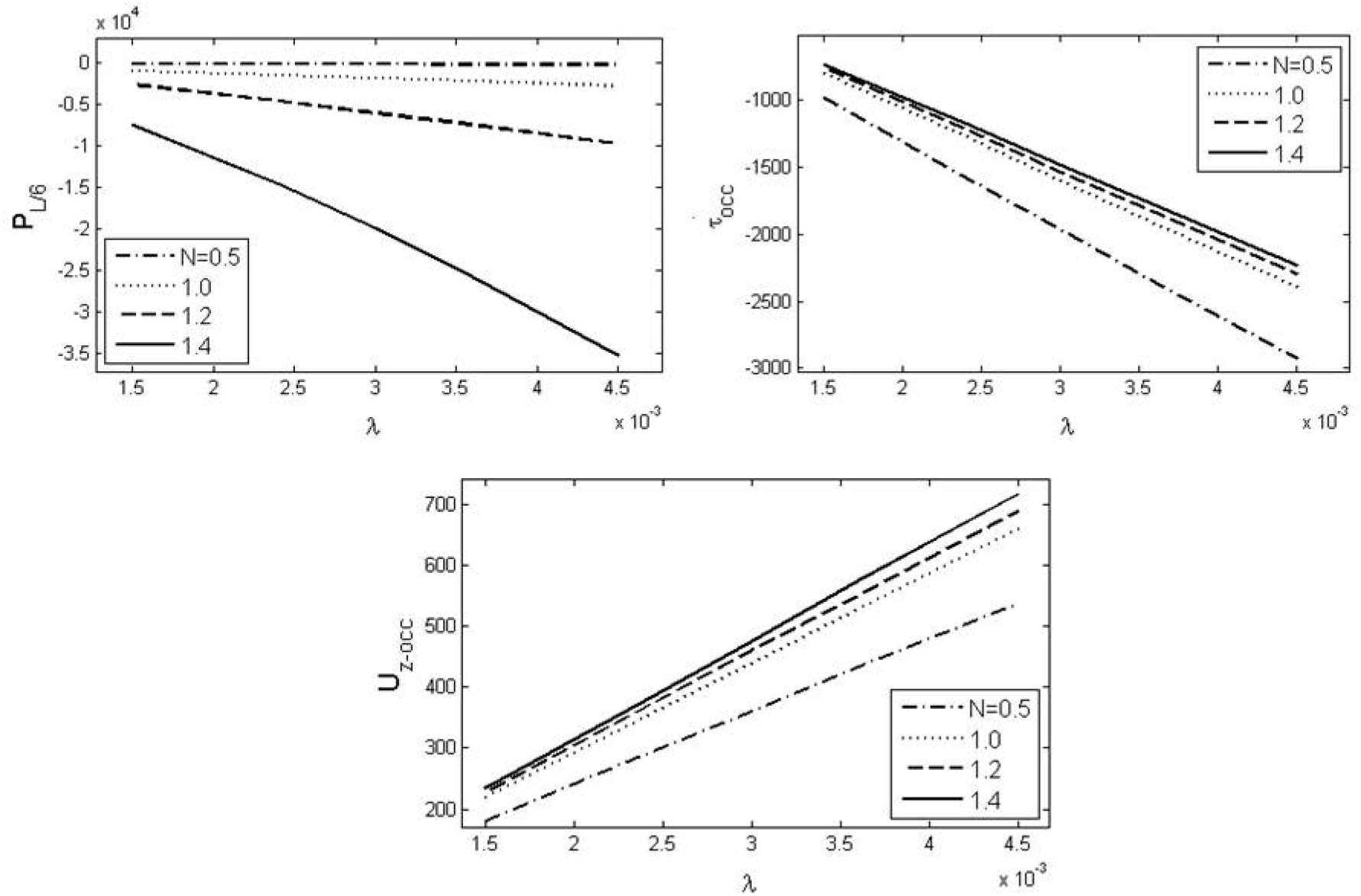
Effect of wavelength on the flow for fluid having flow behavior index = 0.5 (pseudo-plastic), 1.0 (Newtonian), 1.2 (dilatants), and 1.5 (dilatants).

## 5. Conclusion

A mathematical model for a two-indenter driven pump having peristaltic-like movement is developed and studied. We have performed the simulation under the assumptions of low Reynolds number having curvature comparable to the luminal diameter. The indenter was designed to mimic the flow driven by alimentary system of the *C. Elegan*, especially the pharynx. By regulating the motion of the front indenter and fixing the rear indenter at higher occlusion, the flow patterns and pressure variation in the lumen were characterized for various type of fluid modeled using the power law approximation. The study demonstrates following behavior for a two-indenter driven pump (one that closely mimics the pharyngeal activity of the nematode to an extent),

- A large pressure barrier is set up at the rear valve which prevents the fluid to enter into the adjacent compartment and moderately varying pressure at the front valve.
- The opening of lumen (using front value) creates suction in the lumen that drives the inflow of contents from outside to the pharynx.
- Nature of fluid transport is apparently decided by the rate of closure of the front valve and the nature of fluid.
- At higher occlusion the chamber formed in between the two valves are pressurized heavily compared to those of lower occlusion. As a result of this, higher flow rates are developed which has an effect of generating higher fluid stress.
- The pressure generated is highest for the dilatants which are followed by Newtonian and pseudo-plastic fluids.
- Shear stress dependency of frequency is highest for lower values of flow behavior index.

## Acknowledgements

The author received no financial support for the research, authorship, and/orpublication of this article.

## Nomenclature

t: time
r: radial coordinate
θ: aximuthal coordinate
z: axial coordinate
c: velocity scale
f: frequency of closure/opening of the 1st valve
z_0_: reference axial position of the 1st valve
z_0_’: reference axial position of the 2nd valve
a: max. radius of the cylinder
λ: wavelength of the 1st valve
λ’: wavelength of the 2nd valve
R: shape function of cylinder indicating radius
P_occ,max_: max. occlusion of the valves
p_occ_: occlusion of the 1st valve at time ‘t’
p_occ’_: occlusion of the 2nd valve at time ‘t’
u_r_: radial velocity
u_θ_: velocity along th-direction
u_z_: axial velocity
U_r,B_: radial velocity of the wall
m: flow consistency index
n: flow behavior index
μ: viscosity of the fluid
P: pressure
τ: wall shear stress
Q: flow rate
*: indicates dimensional variable

## Abbreviations

C. Elegans: Caenorhabditis elegans
MDDS: Micro Drug Delivery System
μTAS: Micro Total Analysis Systems
POC: Point of Care
PCR: polymerase chain reaction

